# Escalation of Memory Length in Finite Populations

**DOI:** 10.1101/130583

**Authors:** Kyle Harrington, Jordan Pollack

## Abstract

The escalation of complexity is a commonly cited benefit of coevolutionary systems, but computational simulations generally fail to demonstrate this capacity to a satisfactory degree. We draw on a macroevolutionary theory of escalation to develop a set of criteria for coevolutionary systems to exhibit escalation of strategic complexity. By expanding on a previously developed model of the evolution of memory length for cooperative strategies by Kristian Lindgren, we resolve previously observed limitations to the escalation of memory length. We present long-term coevolutionary simulations showing that larger population sizes tend to support greater escalation of complexity than smaller population sizes. Additionally, escalation is sensitive to perturbation during transitions of complexity. In whole, a long-standing counter-argument to the ubiquitous nature of coevolution is resolved, suggesting that the escalation of coevolutionary arms races can be harnessed by computational simulations.

## 1. Coevolutionary Escalation

The escalation of complexity and accretion of knowledge within an evolving population are poorly understood ideas. Yet the study of coevolution and open-ended evolution represents some of the most ambitious research agendas [1] with implications for directed evolution in synthetic biology [2, 3], evolutionary robotics [4], and automatic programming [5]. Long-term evolution studies have been conducted in microbiological systems [6]; however, studies of the evolutionary dynamics of complex strategies in cooperative games have not achieved the same degree of success [7].

The theory of natural selection is historically associated with phylectic gradualism, the slow transformation of one species to another. However, Eldridge and Gould proposed that new species emerge rapidly in punctuated equilibria [8]. These punctuated equilibria are generally associated with an allopatric (geographic) mechanism of species emergence, whereby relocation to a novel environment leads to a change in selection pressure and often a change in population capacity. In this work we show how these innovative evolutionary phenomena can arise solely from coevolutionary interactions, specifically competitive coevolution.

Coevolution describes the dynamics that arise from interactions between species over evolutionary timescales. “Coevolution” was first coined by Ehrlich and Raven as an approach to the study of community evolution [9]. The study of coevolution encompasses many types of community interactions, be they antagonistic, neutral, or symbiotic. A cornerstone of `coevolution is reciprocal selection, where selection on one species reciprocates to other features and members of the ecology. Reciprocal selection has been shown to cause evolutionary arms races in Natural systems [10], where Yucca moth exhibited features indicative of reciprocal adaptation with the Yucca plant. Yet, coevolutionary dynamics have some notable pathologies that make the maintenance of such evolutionary arms races non-trivial.

The literature on computational co-evolution has demonstrated a range of pathologies. Coevolutionary simulations have been plagued by a history of mediocre results and stable states [11, 12]. In one such study the inability of evolutionary game theory to model the dynamics of an evolutionary algorithm with a fitness structure defined for the classic evolutionary hawk-dove game was presented [13]. It was shown that the failure was primarily caused by an insufficient finite population size [14]. This led to the formalization of finite evolutionary stable states within the field of evolutionary game theory [15]. To facilitate the study of coevolutionary pathologies Watson and Pollack developed the Numbers Game [16], which exhibits a range of fundamental coevolutionary pathologies: loss of gradient, focussing, and relativism. Loss of gradient occurs when the fitness with respect to a sample population does not reflect the absolute objective fitness. Focussing occurs when selective pressures focus on a subset of traits, such that the value of other traits can be forgotten. Relativism occurs when selection pressures favor traits of similar quality, relaxing pressures on more advanced traits. The increased rigor in the computational study of coevolutionary dynamics led to the adoption of the game theoretic tool, solution concepts, by the coevolutionary community [17]. Bucci and Pollack then introduced the mathematical framework of maximally informative individuals [18], which resolves a number of coevolutionary pathologies by using a mechanism for ordered sets reminiscent of principle component analysis.

A significant pathology of evolutionary histories is what has become known as the Red Queen effect [19]; a species must adapt as fast as it can just to survive the typical changes of the system. Specifically, after analysis of the fossil record van Valen discovered that the probability of a species’ extinction is generally independent of the age of the species [19]. While the notion of a constant extinction rate has been subject to serious review and is no longer in favor [20], the majority of studies assume a positive non-zero probability of extinction. In the face of a continuous pressure for extinction, how can a population evolve towards higher levels of complexity?

### 1.1 Hypothesis of Escalation

The hypothesis of escalation describes how competition between enemies leads to an increase in complexity and/or investment [21, 22]. The dynamic can be summarized with the following example. Consider an environment with 2 snails, one with a thicker shell than the other, and 1 hungry crab. The crab attempts to consume both snails, but can only break the snail with the weaker shell. The harder shelled snail survives and thus has future chances at reproduction. Unless other selective pressures are applied to the snail (which would be the case in a natural environment), we expect that such an encounter between snails and crabs of successive generations would bias snail morphology toward a harder shell. A similar scenario can be described for 2 crabs of varying strengths and a hard-shell snail. The escalation of the antagonistic traits between these species (shell thickness and crab strength) is familiar from the evolutionary arms race analogy of Dawkins and Krebs [23]. We explore a *reduced* hypothesis of escalation which does not account for geographic distribution, and thus does not permit allopatric speciation. Although this removes one of the primary hypothesized mechanisms of producing punctuated equilibria, genetic variation will still remain a property of our model. We will show that the key observations associated with punctuated equilibria and escalation persist.

The original hypothesis of escalation is a naturalist perspective [22], and details many features of Nature which are suggested as requirements for a coevolutionary system to support the maintenance of evolutionary arms races. The original list of criteria for escalation is concisely recapitulated in [21, 24]. We consider a reduced version of the hypothesis of escalation, where geographic features and [extrinsic events] are disregarded, and populations are unstructured with complete mixing. The criteria for the reduced hypothesis of escalation and their corresponding realizations within this work are:

1. **There must be competition**; each strategy competes against many other strategies.
2. **Competition applies selective pressure**; limited population capacity.
3. **Strategies must be evolvable**; there is always a probability of mutation creating a new individual.

We show how adherence to these criteria allow strategies in a cooperative game to escalate in complexity, exhibiting a coevolutionary arms race.

The reduced hypothesis of escalation that we consider is indeed vastly simplified beyond Vermeij’s original hypothesis. We do not claim that the reduced hypothesis exhibits the same rates of escalation as the original hypothesis, because as Vermeij suggests [21], positive feedback can arise as a result of escalation across a geographic distribution of environments. The hypothesis of escalation has recently be challenged with additional statistical analysis of the fossil record [25]. These analyses have been largely invalidated on the basis of sample selection and the fossilization properties of the studied organisms [26, 27]. There still remains a debate regarding how much of evolutionary history is driven by microevolutionary antagonistic interactions, such as in the case of escalation, and macroevolutionary trends such as punctuated equilibria. We do not attempt to resolve this question, but offer support to the microevolutionary perspective of Vermeij’s hypothesis of escalation. This brings us to our computational model of escalation in a game called the Iterated Prisoner’s Dilemma with noise based upon [7].

## 2. Evolution of Cooperation

The Prisoner’s Dilemma has become the predominant model of the evolution of cooperation. In this game, two players are faced with the choice of deciding to cooperate or defect against their opponent, but their payoff is dependent upon both players’ decisions. Specifically, the best situation for a single player is to defect against a cooperative opponent; the second best situation for a single player (but best for both players combined) is for both players to cooperate. If players have no memory, then the safest assumption is that the other player is rational and will attempt to maximize payoff. Thus, a rational player with no memory will always defect. When the game is extended to multiple rounds of play the game is called the Iterated Prisoner’s Dilemma (IPD), which is the focus of this model. In the IPD a player may decide to cooperate or defect based upon memory of recent encounters with their opponent. For the model presented in this paper, every strategy of a given memory length encodes the response (cooperate or defect) for all possible histories.

In the early 1980’s, Axelrod and Hamilton conducted a computer tournament of human-designed IPD strategies [28]. The winner of the tournament was Anatol Rappaport’s tit-for-tat strategy. Since these initial tournaments a number of researchers have embarked on the quest to find the champion evolutionarily stable strategy IPD strategy. A sequence of findings have shaped the current belief about optimal strategies in the IPD. It was shown that tit-for-tat plays a transitory role in the evolution of IPD strategies [29], and subsequent analysis led to the demonstration of the strength of the win-stay, lose-shift strategy [30]. In the case of the stochastic IPD, where the decision to cooperate or defect is determined by the flip of a genetically biased coin, a recent proof demonstrates the existence of “zero-determinant” (ZD) strategies, where a player can unilaterally specify the payoff received by one’s opponent [31]. This proof marks a significant discovery in the structure of the IPD; however, further research on ZD strategies has revealed that they are not ESS [32]. It has been proven that in alternative formulations of the IPD there are no ESSs [33, 34, 35]. However, these proofs involve features such as discounting of future moves, which are not present in the classic IPD. Recent theoretical work has shown that longer strategies improve the average performance of IPD strategies [36] and that longer memory lengths should evolve over time [37]; however, there have been no empirical studies that show evolutionary trajectories that satisfy this claim.

An innovative study was presented by Lindgren [7] where the set of active IPD strategies change over time, as opposed to most studies of the IPD where only the frequencies of a fixed set of strategies change over time. In Lindgren’s study, strategies evolve by flipping between cooperation or defection as based upon a history of interactions with a memory length measures the number of actions. For example, a memory length of 4 means the strategy is dependent on 2 interactions between both players. However, Lindgren found that the model was not able to escape an ESS containing strategies of memory length 4. In other words, the system did not appear to escalate beyond memory length 4. Our model alleviates this problem by using an alternative variation mechanism. In his model, memory lengths increase by doubling and halving, which only allows for the introduction of mutant strategies that vary by the action of one player from the current population. Instead we introduce mutants with a normal distribution of memory length variations, permitting the mutant strategies of any length. The limitation of changing memory length by only 1 interaction at a time is an inductive bias which expects that a successful mutant exists within a factor of 1 of the current distribution of memory lengths in the population. This intuition can be reinforced by the fact that a memory length extension of 1 only affects a player’s behavior with respect to one role in the game. For example, tit-for-tat is a strategy of memory length 1, where TFT cooperates if its opponent cooperated, and defects if its opponent defected. An extension of memory length 1 allows TFT to remember not only its opponent’s previous move, but also its own previous move. However, we argue that there are situations where a strategy can only be invaded by a mutant who’s strategy has changed by more than 1 memory length. While Lindgren’s operators do not guarantee that the third criteria of the reduced hypothesis of escalation (strategies are improvable by variation) are satisfied, our genetic operators do.

A similar observation on variation was made by Ikegami who uses tree-representations of IPD strategies [38], where his populations exhibit escalation of memory length and diversity. However, Ikegami’s model is obscured by the use of a module-based evolutionary operator. This module-based operator, akin to symbiogenesis, provides a similar variable memory length extension to our normally-distributed extension/contraction operators. However, genetic recombination is an evolutionary transition that is expected to have emerged long after populations began to escalate [39].

It is well known that the size of a finite evolving population can have a significant impact on the fate of the population [40, 14, 15]. However, the relationship between the size of a population and its ability to support the escalation of strategy memory length remains unexplored. This is a particularly significant direction when considering the IPD with noise, which ensures that every element of a strategy has an fitness consequence. We present long-term simulation data that demonstrate a positive correlation between greater population size and the evolution of longer memory lengths, suggesting that increased population size can lead to enhanced evolution of strategic complexity.

## 3. Model

Our model is an extension of Lindgren’s innovative model of the IPD [7], where the use of alternative genetic operators alleviates the mediocre stable-states previously observed. We both suggest that this model satisfies the criteria of the reduced hypothesis of escalation and empirically demonstrate the escalation of complexity in the model.

### 3.1. Prisoner’s Dilemma

The interactions between evolving strategies are specified by the replicator dynamics and the game. We use a standard formulation of the Prisoner’s Dilemma

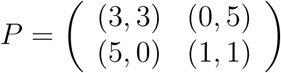

where the notation (*p*_1_, *p*_2_) indicates the scores of players 1 and 2, respectively. However, this payoff matrix only specifies the score of a single round of the Prisoner’s Dilemma. In the Iterated Prisoner’s Dilemma (IPD) multiple rounds are accounted for. The standard way of accomplishing this is by iterating for a finite number of rounds and accumulating the total score for each player during each round. The IPD becomes interesting when strategies have some memory and may change their behavior depending upon the outcomes of previous rounds. This is generally accomplished by encoding lookup-tables within strategies. However, the accumulated score will be sensitive to the number of iterations performed. This will be particularly true as strategies rely on memories of more encounters.

The stochastic Prisoner’s Dilemma admits an alternative method of iteration to the previously mentioned finite iteration technique [41]. Stochastic Prisoner’s Dilemma strategies include a noise term, whereby with a certain probability strategies will take the opposite action. This means the game is a Markov chain. We can describe the game as follows

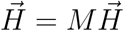

where 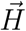 is the vector of probabilities of each history and *M* is the transfer matrix. For two strategies, *s*_1_ and *s*_2_, 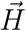 is always of length 2*^max^(|s*_1_*|,|s*_2_*|*). The transfer matrix describes the probabilities of transitioning between histories given *s*_1_ competing against *s*_2_ with noise. *H* is called the stationary distribution of *M*, and represents the distribution of histories in the limit of an infinite number of rounds. We can recover the distribution of round outcomes (CC, CD, DC, and DD) by weighing all histories that end in each outcome by the corresponding payoff values. This distribution of rounds allows us to compute the scores of *s*_1_ and *s*_2_.

### 3.2. Genetic Variation

We employ the same genetic encoding as Lindgren. Strategies are represented as binary strings that encode the action to perform given a specific history. This is easily accomplished by using the observed history as an index into the genome, where the binary value stored at that position specifies the strategy’s response. In the IPD with finite rounds, strategies generally also encode a sequence that specifies the “initial history” because this historical lookup mechanism only works when the genome encodes the responses for all historical sequences. A study of the effect of memory size on the finite-round IPD was presented in [42].

The genetic operators first used by Lindgren [7] implement gene-doubling, gene-halving, and point mutation. Point mutation is familiar from genetic algorithms, where a single bit is flipped with some mutation probability. In gene-doubling, the entire bitstring is extended by a factor of 2 during duplication, because the index is based on the historical observations a doubling event on its own does not change the meaning of a genome. Gene halving is accomplished by randomly truncating the first or second half of the genome.

We use variants of each of these genetic operators. Instead of point mutation, we use uniform mutation, where multiple bits may be flipped during a single reproductive event. To accomplish extension and contraction we draw a random number from a Gaussian distribution, and the absolute value of the integer component is taken as the number of extensions/contractions to perform. Both extension and contraction are accomplished in the same way as Lindgren’s model, but extension/contraction may be more/less than a factor of 2. When performing genetic operations, first the mutant genome may or may not be extended/contracted, then it subsequently may or may not be subject to uniform mutation. Thus in a given reproductive event a mutant may have been extended/contracted as well as varied with uniform mutaiton. We now revisit a requirement of the hypothesis of escalation: “strategies are evolvable.”

Our genetic operators ensure that it is possible to reach a large number of strategies from any population distribution, while Lindgren’s operators appear to only reach a limited range of genotypes. Specifically, no mutant strategy will ever be larger than 1 memory length longer than the biggest genome in the population, or smaller by more than 1 memory length than the smallest genome in the population. However, we hypothesize that it may be necessary to invade with a strategy outside of that range, and our results suggest this is correct. There has been some work on the invasion by pairs of strategies [33, 34, 35], but there is still no known champion IPD strategy.

While Lindgren utilized the continuous-time replicator dynamics and introduced mutants while time-stepping, we instead use the Moran process with mutation. The Moran process models evolutionary dynamics by iteratively replacing one individual at a time [43]. In the evolutionary computation literature the Moran process is sometimes called “steady-state” evolution [44]. The Moran process offers an intuitive way of introducing mutant strategies into the population. On the other hand, the best method of introduction of mutant strategies into a mean-field model is not immediately apparent. In Lindgren’s model, each strategy has a probability of introducing a single mutated variant proportional to the frequency of the parent strategy.

## 4. Results

*Average diversity per population size over time.* In this study we simulate the previously described model with the Moran process using a range of population sizes. For all simulations the following parameters are used: *p_extend_* = 0.000001, *p_contract_* = 0.000001, *p_uniform_* = 0.001, and *T_max_* = 100, 000 generations. We have also restricted the maximum length of strategies to 12; however, we never observe this limit being reached. The cost of simulating infinite games increases exponentially with the maximum memory size of the competing strategies, which is a strong motivation for prohibiting excessively long strategies.

Just as in Lindgren’s study [7], we observe similarities between all simulations (and Lindgren’s), especially during the initial generations as the system passes through meta-stable states. For example compare Figure 1 to Lindgren’s Figure 1 [7], both of which exhibit the same patterns in the initial phase of their evolutionary trajectory. While much of Lindgren’s discussion regarding the evolutionary timeline remains intact, our model provides an epilogue to Lindgren’s allusion to open-ended evolution.

**Figure 1:**
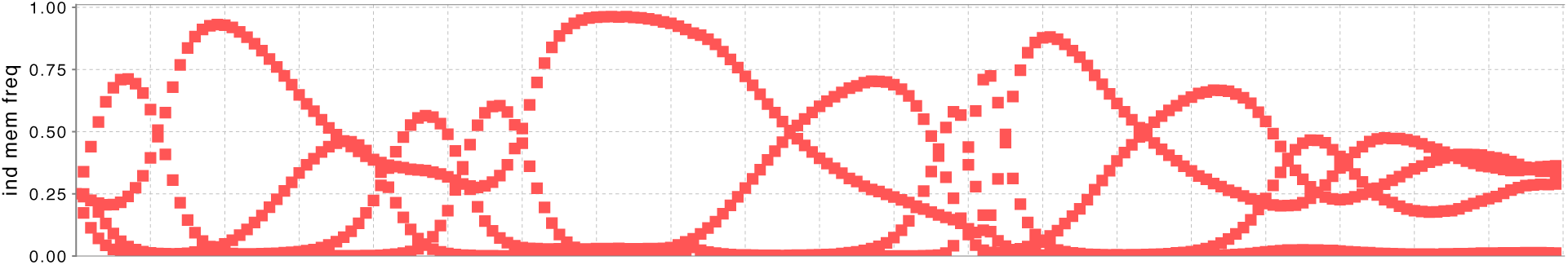
Example of initial evolution.

The inclusion of noise in the IPD model admits evolutionarily stable strategies [45]. Both Lindgren’s and our model do reach evolutionarily stable states under some conditions, and in Lindgren’s model it is unclear whether all paths will lead to such an ESS. While Lindgren found some evolutionary trajectories that did not get stuck in the same memory 4 ESS that plagued many of his simulations, he did not demonstrate evolutionary trajectories that exceeded memory length 4. Here we present simulation results for evolutionary trajectories that escalate beyond memory length 4.

We conducted experiments using the Moran process with population sizes: 5,000 and 10,000. 25 replicates were used for each population size. Although we were not able to simulate all population sizes for the full 100,000 generations, we present results where a number of evolutionary trajectories pass the memory 4 meta-stable state. When comparing results each timestep represents a generation, which is *N* breeding events, where *N* is the population size. Therefore, when 2 simulations are compared with different population sizes, at any given timestep each population will have experienced a different number of reproductive events (i.e. *N* ∗ *t*).

The smallest population size that we consider is 5,000 (Figure 3A). In this case a number of simulations are unable to escape the initial meta-stable states, and populations remain at low memory lengths. However, some simulations do reach populations consisting of primarily memory length 7. We will revisit this observation for some of a larger population size. The number of species grows for the first 70,000 generations, then plateaus just below 300 species. However, even during this plateau of species diversity, some escalation can still be observed as strategies of memory length 7 are still on the rise at generation 100,000.

**Figure 2:**
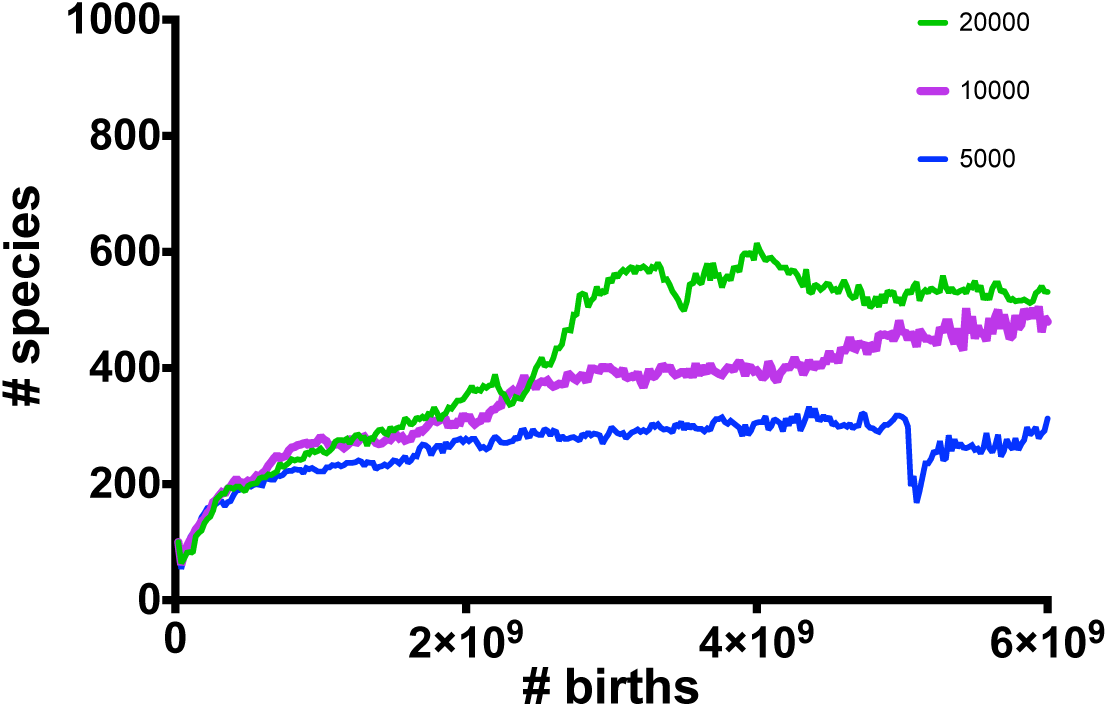
Species diversity for population sizes 5,000, 10,000, and 20,000 averaged over 25 simulations.

**Figure 3:**
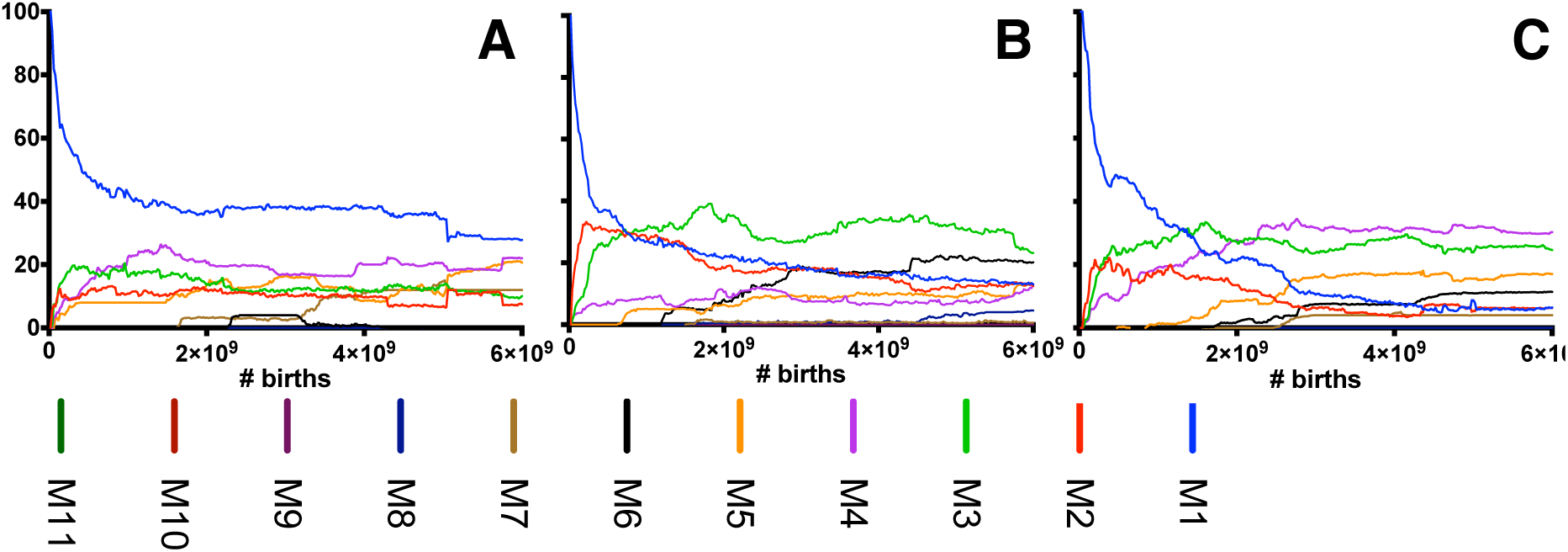
Fraction of population composed of each memory length for Population size 5,000, 10,000, 20,000. Runs are averaged over 25 simulations.

Our results for population sizes 10,000 and 20,000 show the most escalation in memory length (Figures 3B-C). Results the population size of 20,000 show runs that are beginning to be dominated by memory length 8. These runs are the most escalated of all experiments we conducted. The average number of species is similar across the runs to the extent that the runs are completed. However, the correlation between the completion of runs and number of species is clear. Spikes in the number of species significantly slow simulation speeds. For this reason, we cannot make clear statements regarding the number of species supported by each population size.

By extending Lindgren’s model with alternative genetic operators we have cleared the path to open-ended evolution in the IPD model. We explore the model using finite population evolutionary dynamics, as opposed to Lindgren’s use of continuous-time replicator dynamics. The model continues to exhibit similar evolutionary trajectories to those presented by Lindgren, which suggests that it is not our use of the Moran process that leads to the escape from the memory 4 meta-stable states that appeared to limit Lindgren’s original model. The computational cost of simulating large population sizes causes us to present partial results. While we see that larger population sizes are capable of supporting a larger number of species, larger population size does not eliminate the possibility of getting stuck in an evolutionary equilibrium. This leads to the suggestion that achieving greater escalation is not simply a matter of using a larger population size.

Now let us consider a specific example trajectory from a population size 20,000 run. In Figure 4 we have a timeline showing the evolutionary history after 2 *∗* 10^9^ birth events. Over the course of this evolutionary trajectory the population transitions to the previously observed limit of memory length 4 to memory length 6 and on to memory length 8. As we noted in Figure 2, the diversity of species increases significantly toward the end of population making analysis of individual strategies challenging. To this end we perform a species knockout analysis at multiple points within the evolutionary trajectory.

**Figure 4:**
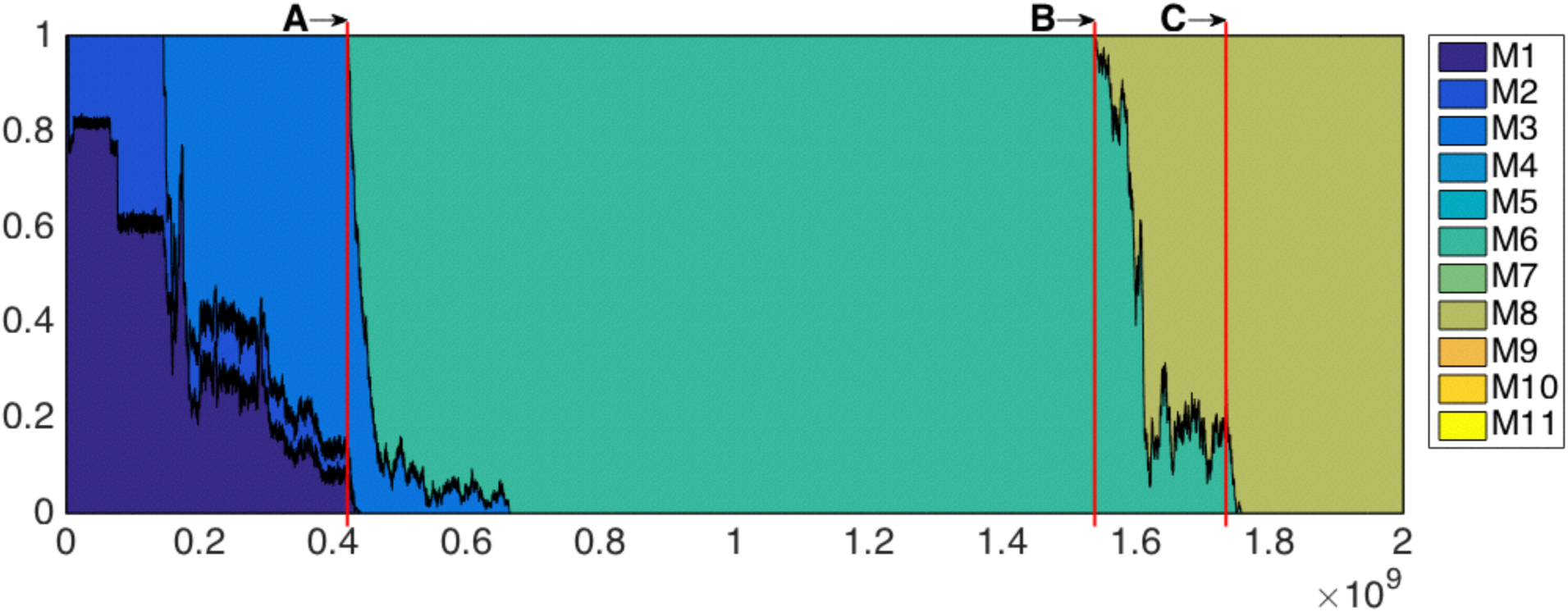
Fraction of population composed of each memory length for an example run with population size of 20,000. Annotated vertical lines indicate populations where specie knockout analyses are performed.

*Species knockout analysis.* An analysis of population stability was performed via species knockout, where a given strategy is eliminated from the population. The population is rebalanced by uniformly allocating the previously occupied fraction of the population to the remaining strategies. After performing the knockout, the simulation is evaluated for 4 *×* 10^5^ birth events, then the distribution of memory lengths are investigated. For each timepoint a species knockout is performed with respect to each strategy in the population. The timepoints of knockouts are denoted in Figure 4.

Knockout A is performed at the transition from memory length 4 to 6. Memory length 4 is the level where Lindgren found a tendency for populations to stabilize [7]. The knockout is performed at the first generation where memory length 6 strategies appear. Of the 17 species knockouts performed, 2 knockouts lead to a collapse of escalation, while the remaining knockouts continue to support memory length 6 strategies.

Knockout B is performed at the first timepoint where there are more than 50 individuals with memory length 8 strategies. It was necessary to choose such a timepoint because mutations ephemerally introduce strategies of memory length 8 that are not capable of triggering a transition to memory length 8. Nevertheless, for all 41 knockouts the populations revert to memory length 6. This suggests that the fitness of strategies are highly interdependent during this particular transition to greater complexity, which motivates us to consider a knockout after the transition from memory length 6 to 8 has progressed further.

Knockout C is performed when the majority of the population is occupied by memory length 8 strategies (approx 30% to 70%). Here we find that all 157 knockouts maintain populations with memory length 8 strategies. This suggests that the interdependence observed in knockout B has stabilized, and the population has become more robust to the distribution of strategies that it contains.

These three knockout studies highlight a key point. Knockout A is performed immediately following the transition from memory length 4 to 6 and still many knockout populations are capable of escalating to greater memory lengths. Knockout B is performed close to the transition from memory length 6 to 8 and none of the knockout populations escalate to greater memory lengths. Finally, knockout C is performed much later in the transition from memory length 6 to 8 and all knockout populations continue to escalate to greater memory length. While it is possible that longer evaluation of knockout populations may lead to observations of eventual escalation to greater memory length, in this example the point remains that that escalation of greater complexity is more vulnerable to destabilizing knockouts.

## 5. Conclusion

The study of coevolutionary arms races has had a challenging history plagued with premature mediocre stabilization [12] and other coevolutionary pathologies [16]. These pathologies were previously related to observations of limited escalation of complexity in simple evolutionary models of cooperative games [7]. In our study we have drawn inspiration from the macroevolutionary theory of the escalation of coevolutionary interactions [22] to show that previous observations of limited evolution in the Iterated Prisoner’s Dilemma with noise [7] were due to a lack of evolvability. By conducting long-term evolutionary simulations we have shown that an improved model can lead to continued escalation of strategic complexity. We have also shown that strategies escalate in complexity faster in larger populations. Coevolutionary simulation can drive the escalation of complexity and that escalation can be amplified in larger population sizes. Furthermore, the escalation of complexity can be sensitive to species knockouts during transition periods. Thus, the stabilization of species and maintenance of large population sizes are viable mechanisms to supporting the escalation of strategic complexity.

## Acknowledgements

We thank Sevan Ficici and Anthony Bucci for insightful discussions, and Kristian Lindgren for sharing his code.

